# Golgi anti-apoptotic proteins are evolutionarily conserved ion channels that regulate cell death in plants

**DOI:** 10.1101/859678

**Authors:** Maija Sierla, David L. Prole, Nuno Saraiva, Guia Carrara, Natalia Dinischiotu, Aleksia Vaattovaara, Michael Wrzaczek, Colin W. Taylor, Geoffrey L. Smith, Bart Feys

## Abstract

Programmed cell death regulates developmental and stress responses in eukaryotes. Golgi anti-apoptotic proteins (GAAPs) are evolutionarily conserved cell death regulators. Human and viral GAAPs inhibit apoptosis and modulate intracellular Ca^2+^ fluxes, and viral GAAPs form cation-selective channels. Although most mammalian cell death regulators are not conserved at the sequence level in plants, the GAAP gene family shows expansion, with five paralogues (*AtGAAP1-5*) in the Arabidopsis genome. We pursued molecular and physiological characterization of AtGAAPs making use of the advanced knowledge of their human and viral counterparts. Structural modeling of AtGAAPs predicted the presence of a channel-like pore, and electrophysiological recordings from purified AtGAAP3 reconstituted into lipid bilayers confirmed that plant GAAPs can function as ion channels. AtGAAP1 and AtGAAP4 localized exclusively to the Golgi within the plant cell, while AtGAAP2, AtGAAP3 and AtGAAP5 also showed tonoplastic localization. Gene expression analysis revealed differential spatial expression and abundance of transcript for *AtGAAP* paralogues in Arabidopsis tissues. We demonstrate that AtGAAP1-5 inhibit Bax-induced cell death in yeast. However, overexpression of AtGAAP1 induces cell death in *Nicotiana benthamiana* leaves and lesion mimic phenotype in Arabidopsis. We propose that AtGAAPs function as Golgi-localized ion channels that regulate cell death by affecting ionic homeostasis within the cell.

**Highlight:** Arabidopsis Golgi anti-apoptotic proteins (GAAPs) share functional conservation with their human and viral counterparts in cell death regulation and ion channel activity

**Abbreviations:** AtGAAP, *Arabidopsis thaliana* GAAP; BI-1, Bax inhibitor-1; CFP, cyan fluorescent protein; CMLV, camelpox virus; ER, Endoplasmic reticulum; GAAP, Golgi anti-apoptotic protein; GFP, green fluorescent protein; hGAAP, human GAAP; LFG, Lifeguard; LMM, lesion mimic mutant; PCD, programmed cell death; TMBIM, transmembrane Bax inhibitor-1 motif-containing; TMDs, transmembrane domains; vGAAP, viral GAAP; YFP, yellow fluorescent protein

## INTRODUCTION

Programmed cell death (PCD) is a process that plays essential roles in the life cycle of plants and animals. In multicellular organisms, PCD is a key mechanism controlling developmental pattern formation, organ shape, and the removal of unwanted, damaged or infected cells. In plants, developmental PCD contributes to many processes, such as death of suspensor cells in the developing embryo (Bozhkov *et al*., 2005), tracheary element differentiation (Fukuda, 2000) and organ senescence (Thomas, 2013). In addition to its role in development, PCD in plants can also be induced by abiotic and biotic cues. Rapid induction of PCD at the site of pathogen infection, the hypersensitive response (HR), is a powerful defense mechanism that limits the spread of biotrophic pathogens in plants (Coll *et al*., 2011). A variety of abiotic stresses, including heat-shock (Vacca *et al*., 2004; Watanabe and Lam, 2006), low temperature (Koukalova *et al*., 1997), ultraviolet irradiation (Danon *et al*., 2004) and ozone exposure (Overmyer *et al*., 2005) also induce PCD.

Despite its central importance, the molecular mechanisms governing the initiation and execution of PCD in plants are largely unknown. This contrasts with animals, where apoptosis is a well characterized form of PCD (Dickman *et al*., 2017). Mitochondrial pathway of apoptosis involves release of cytochrome c from mitochondria into the cytoplasm, which triggers a signaling cascade leading to activation of caspases, a family of cysteine proteases, and subsequently apoptosis. Bcl-2 proteins are core regulators of apoptosis that can either inhibit (e.g. Bcl-2 and Bcl-XL) or promote (e.g. Bax and Bak) apoptosis by controlling the integrity of the mitochondrial outer membrane (Chipuk *et al*., 2010).

PCD in animals and plants shares several morphological and biochemical features, including cell shrinkage, DNA fragmentation, Ca^2+^ fluxes, and production of reactive oxygen species (ROS) (Dickman *et al*., 2017). No direct homologues of mammalian caspases have been identified in plants, but caspase-like protease activities are associated with the activation of various types of PCD in plants (Coffeen and Wolpert, 2004; Hatsugai *et al*., 2004; Kuroyanagi *et al*., 2005; Hatsugai *et al*., 2006; Hatsugai *et al*., 2009). Similarly, although plants have no apparent homologues of Bcl-2 proteins, expression of mammalian Bax triggers PCD in Arabidopsis (*Arabidopsis thaliana*) and tobacco (Lacomme and Santa Cruz, 1999; Kawai-Yamada *et al*., 2001). Although much of the core machinery of animal apoptosis is not conserved in plants at the sequence level, two related families of anti-apoptotic proteins are present in both animals and plants: Bax inhibitor-1 (BI-1) and Golgi anti-apoptotic protein (GAAP). BI-1 proteins were originally identified as inhibitors of Bax-induced cell death in yeast (Xu and Reed, 1998), and their roles in the regulation of cell death in both animal and plant cells are now established. BI-1 inhibits cell death by controlling the Ca^2+^ content of the endoplasmic reticulum (ER) and intracellular Ca^2+^ fluxes (Ishikawa *et al*., 2011; Robinson *et al*., 2011).

GAAPs are an evolutionarily conserved group of anti-apoptotic proteins that were originally discovered in poxviruses (Gubser *et al*., 2007; Carrara *et al*., 2017). GAAP orthologues have been identified throughout eukaryotes, including animals, plants and fungi, as well as some prokaryotes (Gubser *et al*., 2007; Carrara *et al*., 2015; Carrara *et al*., 2017). Over-expressing human (hGAAP) or viral (vGAAP) GAAP in human cells protects against both intrinsic (e.g. Bax) and extrinsic (e.g. Fas) pro-apoptotic stimuli. Silencing of hGAAP with small interfering RNA (siRNA) leads to cell death in human cells, suggesting that hGAAP is essential for cell viability (Gubser *et al*., 2007). vGAAP can complement for the loss of hGAAP, indicating functional conservation (Gubser *et al*., 2007). Cell adhesion, spread and migration are also regulated by hGAAP (Saraiva *et al*., 2013a). In mammalian cells, over-expression of hGAAP or vGAAP reduces the Ca^2+^ content of the Golgi and ER, while knock-down of hGAAP has the opposite effects (de Mattia *et al*., 2009; Saraiva *et al*., 2013a; Saraiva *et al*., 2013b). The increases in cytosolic and mitochondrial Ca^2+^ concentration evoked by an apoptotic stimulus (staurosporine) and by IP_3_-evoked Ca^2+^ release from intracellular stores are reduced by over-expression of hGAAP (de Mattia *et al*., 2009). Purified vGAAP and human BI-1 form cation channels in lipid bilayers (Carrara *et al*., 2015), suggesting that they may directly mediate a Ca^2+^ leak from the Golgi and ER, thus reducing the amount of Ca^2+^ available for release in response to apoptotic stimulation. Since intracellular Ca^2+^ fluxes affect the sensitivity of cells to apoptosis (Giorgi *et al*., 2012), hGAAP might suppress apoptosis by down-regulating cytosolic and mitochondrial Ca^2+^ signals (de Mattia *et al*., 2009).

The molecular and physiological functions of plant GAAPs remain largely uncharacterised. Roles for plant GAAPs have been proposed in plant-fungal interactions (GAAP referred to as Lifeguard (LFG); Weis *et al*., 2013), ER stress responses (Wang *et al*., 2019; Guo *et al*., 2018) and brassinosteroid signaling (Yamagami *et al*., 2009). However, it remains unexplored whether plant GAAPs share functional conservation in cell death regulation and ion channel properties with their animal and viral counterparts. Here, we show that an expanded family of GAAPs exists in plants and undertake a thorough functional characterization of the five Arabidopsis GAAPs (AtGAAPs). Homology modeling was used to predict the structures of AtGAAPs and define a putative ion channel pore. Experimentation showed that AtGAAPs form ion channels in lipid bilayers and inhibit Bax-induced cell death in yeast. Fluorescently-tagged AtGAAPs displayed subtype-specific localization within the Golgi and tonoplast of cells *in planta*, and gene expression analyses show that the five AtGAAP display different tissue distributions in Arabidopsis, suggesting subtype-specific functional roles of AtGAAPs. Arabidopsis strains with single, double and triple knockouts of AtGAAPs did not display obvious morphological phenotypes, which may suggest a high degree of redundancy in the function of AtGAAPs, or that AtGAAPs regulate responses to specific environmental stimuli or stresses. In contrast, fluorescently-tagged AtGAAP1 induced cell death when over-expressed *in planta*, demonstrating a role for AtGAAPs in the regulation of cell death in Arabidopsis.

## MATERIALS AND METHODS

### Protein identification and phylogenetic analysis

Plant GAAP and BI-1 sequences were retrieved from the National Center for Biotechnology Information (NCBI; https://www.ncbi.nlm.nih.gov/) and Phytozome (https://phytozome.jgi.doe.gov) (Goodstein *et al*., 2012) databases using hGAAP and hBI-1 sequences as initial queries, followed by queries using identified Arabidopsis sequences. Amino acid sequences were aligned using Pagan (Löytynoja *et al*., 2012) and the Maximum-Likelihood tree was constructed in RAxML (Stamatakis, 2014) with 1000 bootstrap replicates. Percentage identity and similarity of sequences were calculated using Basic Local Alignment Search Tool (BLAST) (Altschul and Lipman, 1990).

### Sequence alignments and topology predictions

Sequence alignments were generated using Clustal Omega (European Molecular Biology Laboratory-European Bioinformatics Institute) (Sievers *et al*., 2011). The GenBank accession numbers for the sequences used were: NP_193209 (AtGAAP1); NP_191890 (AtGAAP2); NP_192178 (AtGAAP3); NP_567466 (AtGAAP4); NP_171806 (AtGAAP5); O31539 (BsYetJ) and AAG37461 (camelpox virus, CMLV, GAAP). TMDs were predicted using TOPCONS (Bernsel *et al*., 2009). Asterisks indicate positions of fully conserved residue, while colons indicate residues with highly similar properties.

### Structural modelling

Structural modelling was performed as described (Carrara *et al*., 2015). I-TASSER (Zhang, 2008; Roy *et al*., 2010; Roy *et al*., 2012) was used to create homology models of AtGAAPs. The sequences of AtGAAPs were used in structure-based sequence alignments that search for suitable structures within the Protein Data Bank (PDB). The sequences used were: NP_193209 (AtGAAP1); NP_191890 (AtGAAP2); NP_192178 (AtGAAP3); NP_567466 (AtGAAP4); and NP_171806 (AtGAAP5). The searches showed that BsYetJ structures as templates gave the best models. The crystal structures of BsYetJ in closed (PDB: 4PGR) and open (PDB: 4PGS) states were used as templates for the models shown (Chang *et al*., 2014). These models had confidence scores (*C*-scores) in the range −1.21 to +0.06, which are indicative of correct models (Roy *et al*., 2010).

### Protein expression and purification in yeast

Protein expression and purification was carried out as reported (Carrara *et al*., 2015). AtGAAP3 was expressed in *S. cerevisiae* strain FGY217 (Kota *et al*., 2007), under control of the galactose promoter (Mumberg *et al*., 1995). Recombinant AtGAAP3 was purified in 150 mM NaCl, 20 mM Tris-base, 5% glycerol and 0.06% lauryldimethylamine N-oxide (LDAO), pH 7.5 and analysed according to a protocol developed for transmembrane proteins (Drew *et al*., 2008). Briefly, the GFP-8His tag was cleaved by adding 8His-tagged tobacco etch virus (TEV) protease to the purified GFP-8His-tagged AtGAAP3 at a molar ratio of 1:1, and digested overnight at 4°C. Cleaved GFP-8His and the His-tagged protease were removed using a HisTrap nickel column (GE Healthcare), and the untagged target AtGAAP3 was harvested from the flow-through. The purified protein was concentrated using an Amicon Ultra centrifugal filter with a molecular mass cut-off (MWCO) of 30 kDa (Millipore) and analysed on a Superdex 200 size-exclusion chromatography (SEC) column (GE Healthcare). Target protein fractions were collected and concentrated to 1.5 - 2 mg/ml.

### Reconstitution of proteins into giant unilamellar vesicles (GUVs)

GUVs were produced as reported (Carrara *et al*., 2015) by electroformation from a mixture of 1:10 cholesterol (Sigma) to 1,2-diphytanoyl-*sn*-glycero-3-phosphocholine (DPhPC) (Avanti Polar Lipids) dissolved in chloroform (Carl Roth). The lipid mixture (20 μl) was spread on the conductive side of an indium tin oxide (ITO)-coated slide, and dried for 10 min. The dried lipid film was covered with 1 M sorbitol (250 μl), enclosed within a greased O-ring and overlaid with another ITO-coated slide with its conductive side facing the lipids. The assembly was connected to a Vesicle Prep Pro (Nanion). Parameters of the electric field used for electroformation of GUVs were: 5 Hz, 3 V, for 128 min at 20°C. GUVs were re-suspended from the slide within the sorbitol overlay, collected and stored for 3-4 days at 4°C. Proteins were incorporated into GUVs as reported (Carrara *et al*., 2015) by mixing purified protein (10 μl) with GUVs (90 μl) (0.2 mg/ml final protein concentration). Bio-Beads SM-2 absorbents (152-8920, Bio-Rad) that had been washed in methanol (3 x 10 min), ethanol (3 x 10 min) and Milli-Q water (6 x 5 min) were added (40 mg/ml, 15 min), and then removed three times during the reconstitution procedure (45 min total incubation) to remove excess detergent micelles. GUVs containing protein were stored at 4°C and used within a few h for recordings.

### Electrophysiological recording

Single-channel recordings were performed as reported (Carrara *et al*., 2015) with a Port-a-Patch system (Nanion) (Fertig *et al*., 2002; Bruggemann *et al*., 2003), using NPC-1 borosilicate glass chips (5-10 MΩ resistance). GUVs in 1 M sorbitol (5 μl) were added to the *cis* side of the chip and planar lipid bilayers were formed across the aperture using suction (see Figure 3D). Bilayer formation generated seal resistances of 1-10 GΩ. Protein-reconstituted GUVs (5 μl) (in 15 mM NaCl, 2 mM Tris-base, 0.5% glycerol, 0.006% LDAO and 0.9 M sorbitol, pH 7.25) were then added to allow incorporation of purified protein into the bilayer. The final composition of the medium in the *trans* chamber (5 μl) was: 140 mM KCl, 200 nM free Ca^2+^ (220 μM CaCl_2_ buffered with 0.5 mM BAPTA-Na_4_), 10 mM HEPES-free acid, adjusted to pH 7 with KOH. In the *cis* chamber (15 μl, ground) the final composition of the medium was: 46.7 mM KCl, 200 nM free Ca^2+^ (73 μM CaCl_2_ buffered with 0.17 mM BAPTA-Na_4_), 5 mM NaCl, 3.33 mM HEPES-free acid, 0.67 M sorbitol, 0.67 mM Tris-base, pH 7 (see Figure 3D). Recordings were acquired in the “on cell” mode using the PatchMaster software (Nanion) and an EPC 10 patch-clamp amplifier (HEKA). Voltages are expressed as the potential on the *cis* side relative to the *trans* side. Single-channel currents are shown such that downward deflections represent cations flowing from the *trans* to the *cis* side of the bilayer. Continuous current recordings were acquired by applying holding potentials for 1 min durations in increments of 20 mV, or until bursts of spontaneous channel activity appeared. Data were filtered at 2.9 kHz (Bessel filter, HEKA amplifier), digitized at 50 kHz and exported to Clampfit (Molecular Devices) via MatLab (MathWorks). Recordings were analysed with PatchMaster and Clampfit software.

### Cloning and recombination procedure for generation of yeast expression constructs

The coding region of Bax from pBM272 Bax (Addgene) was amplified by PCR using forward (5’-caccaataatggatgggtccggggagcagc) and reverse (5’-tcagcccatcttcttccagatggtgagc) primers, and inserted into pENTR/D-TOPO using a pENTR/D-TOPO Cloning Kit (Invitrogen). Gateway LR Clonase (Invitrogen) was then used to transfer the coding region of Bax into pAG303GAL-ccdb or pAG305GAL-ccdb vectors (Addgene) to give pAG303GAL-Bax and pAG305GAL-Bax. Green fluorescent protein (GFP) from the Gateway entry vector pENTR4-GFP-C1 (Addgene) was transferred to pYES-DEST52 and pAG426GAL-ccdb using Gateway LR Clonase, to create pYES-GFP and pAG426GAL-GFP. Yellow fluorescent protein (YFP) was transferred from a Gateway entry vector to pYES-DEST52 using Gateway LR Clonase, to form pYES-YFP. Gateway LR Clonase was used to transfer AtGAAP, AtGAAP-GFP and AtGAAP-YFP inserts from Gateway entry vectors to pYES-DEST52 (Invitrogen) and pAG426GAL-ccdb (Addgene). Sequences and orientations of all constructs were verified by sequencing (Source Bioscience).

### Yeast apoptosis assays

Strains of *S. cerevisiae* expressing two chromosomally integrated copies of either Bax (termed Bax_2_) or control GFP (termed GFP_2_) under control of the galactose-inducible GAL1 promoter were constructed as follows. The isogenic BY4742 strain of *S. cerevisiae* (MATα, his3Δ1, leu2Δ0, lys2Δ0, ura3Δ0; Euroscarf) was transformed with pAG303GAL-Bax or pAG303GAL-GFP using a Frozen-EZ yeast transformation kit (Zymo Research). Transformants (termed Bax_1_ and GFP_1_, respectively) were selected on agar containing synthetic defined (SD) medium with glucose and dropout supplement lacking histidine (SDglucose-His). Bax_1_ and GFP_1_ yeast were then cultured in SDglucose-His medium at 30°C and transformed with pAG305GAL-Bax or pAG305GAL-GFP, respectively. The resulting transformants (Bax_2_ and GFP_2_, respectively) were isolated on agar containing SDglucose with dropout supplement lacking histidine and leucine (SDglucose-His-Leu). To assess the effects of untagged GAAPs on apoptosis, the Bax_2_ strain was transformed with GFP, AtGAAPs or AtBI-1 in the vector pYES-DEST52, while the GFP_2_ strain was transformed with GFP in pYES-DEST52. For assessing the effects of fluorescently-tagged GAAPs on apoptosis, the Bax_2_ strain was transformed with pAG426GAL-GAAP-GFP, pAG426GAL-GAAP-YFP, pAG426GAL-GFP or pAG426GAL-YFP, while the GFP_2_ strain was transformed with pAG426GAL-GFP. Transformants were selected on agar containing SDglucose with dropout supplement lacking histidine, leucine and uracil (SDglucose-His-Leu-Ura). Colonies were then grown overnight at 30°C in SDglucose-His-Leu-Ura. Cultures were adjusted to identical OD_600_ and a 10-fold dilution series was spotted on either SDglucose-His-Leu-Ura or SDgalactose-His-Leu-Ura selective agar plates (7 µl of each dilution). Growth at 30°C is shown after 2 days (glucose), 6 days (galactose), or 10 days (galactose) for the slowest growing strains. Results are representative of experiments from three independent transformations. Growth of yeast was quantified by measuring the mean pixel intensity of spots (for the 10^1^ dilution) after background subtraction (ImageJ) and normalizing to the mean pixel intensity of spots formed by the GFP_2_+GFP strain on SDgalactose-His-Leu-Ura agar (10^1^ dilution). Media and dropout supplements for the growth of yeast were obtained from ForMedium.

### Plant materials and growth conditions

Both wild-type (Columbia-0 and Landsberg *erecta*) and mutant Arabidopsis plants were grown in a controlled growth room under 16 h light/ 8 h dark (referred to as long day) or 10 h light/ 14 h dark (referred to as short day) cycles, approximately 60% humidity, light intensity of 130 μE m^-2^ s^-1^ and at a temperature of 23°C. *Nicotiana benthamiana* plants were grown under 16 h light/ 8 h dark cycles and under light intensity of 200 μE m^-2^ s^-1^, humidity and temperature as above. Soil-grown plants were sown on seed and modular compost plus sand (Levington) and germinated under short day conditions for two weeks after which plants were transferred to fresh soil. Plants were then placed under long day conditions to induce flowering when necessary.

### Cloning and recombination procedure for generation of transgenic plants

PCR fragments were amplified by PCR using Phusion High-Fidelity DNA Polymerase (Finnzymes) using gene-specific primers. Col-0 genomic DNA was extracted using DNeasy plant mini kits (Qiagen) and used as template for the PCR. PCR products were cloned into the pENTR/D-TOPO vector following manufacturer’s instructions (Thermo Fisher). Inserts were sequenced and recombined into Gateway destination vectors by LR recombination using Gateway LR ClonaseII Enzyme Mix (Thermo Fisher). The pGWB3 destination vector was used to create promoter-GUS fusions. The pGWB5 and pEG101 vectors were used for cauliflower mosaic virus 35S promoter-driven expression of AtGAAPs, with C-terminal GFP tags or YFP tags, respectively. The pGWB vector series was a kind gift from Dr Tsuyoshi Nakagawa (Research Institute of Molecular Genetics, Matsue, Japan) (Nakagawa *et al*., 2009) and the pEarleyGate vectors were described (Earley *et al*., 2006). The AtGAAP and empty vector constructs were transformed into *Agrobacterium tumefaciens* GV3101 cells by electroporation. Electroporation was performed in a pre-chilled 2 mm cuvette, using a BioRad Gene Pulser according to the manufacturer’s recommendations (capacitance 25 μF, resistance 200 Ω, voltage 1.8 kV). Cells were then transferred to 1 ml of SOB medium (2% Difco Bacto-tryptone, 0.5% yeast extract, 0.05% NaCl, 2.5 mM KCl, 10 mM MgCl_2_, and 10 mM MgSO_4_) and incubated for 40 min at 28°C incubator. The transformed cells were selected on LB plates supplemented with the following antibiotics: 50 μg/ml rifampicin, 50 μg/ml gentamycin, 50 μg/ml hygromycin and 50 μg/ml kanamycin for pGBW vectors; 50 μg/ml rifampicin, 50 μg/ml gentamycin and 50 μg/ml kanamycin for pEG vectors. Colony PCR was performed to confirm the presence of the insert with gene-specific primers. Individual colonies were resuspended in 15 μl of sterile water and 2 μl of this was used subsequently as a template in PCRs.

### Arabidopsis transformation and selection of transgenic plants

Binary vectors were transformed into Arabidopsis plants by floral dip (Clough and Bent, 1998). The pGWB3 and pGWB5 transgenic seeds were selected on 0.5 x Murashige and Skoog (MS) plates containing 50 μg/ml hygromycin B. The pEG101 seeds were grown on soil for seven days and screened for Basta (glufosinate-ammonium) resistance.

### Reverse transcriptase PCR (RT-PCR)

Total RNA was isolated using TRI REAGENT (Sigma) according to manufacturer’s instructions and quantified using a spectrophotometer (BioPhotometer, Eppendorf). RNA (2 µg) and an anchored oligo(dT) primer were used for cDNA synthesis using SuperscriptTM III Reverse Transcriptase (Invitrogen) as instructed. cDNA was used as template for PCR with gene specific primers using Phusion High-Fidelity DNA Polymerase (Finnzymes) according to manufacturer’s instructions.

### GUS staining

Plant tissue was immersed in GUS staining solution: 1 mM X-Gluc, 0.1% (w/v) Triton X-100, 0.5 mM K_3_Fe(CN)_6_ (ferricyanide), 0.5 mM K_4_Fe(CN)_6_•3H_2_0 (ferrocyanide), 10 mM Na_2_EDTA, 50 mM PO_4_ buffer, pH 7.0. Tissue was infiltrated twice for two mins in a vacuum chamber before placing samples at 37°C for 16 h. GUS solution was then removed and the tissue was washed twice with 70% ethanol.

### Lactophenol trypan blue staining

Visualisation of dead and dying cells was performed as described (Koch and Slusarenko, 1990). Leaves or leaf discs were collected and immersed in staining solution (25% v/v of each H_2_0, lactic acid, phenol and glycerol plus 0.025% w/v trypan blue) and boiled for 5 mins. Staining solution, once cooled down, was replaced by de-staining solution (250% w/v chloral hydrate). Samples were shaken for 7 days until leaves were cleared. Leaves were mounted in 60% glycerol and viewed with a Leica light microscope (MZ16F, Leica Microsystems).

### Expression of fusion proteins in *Nicotiana benthamiana*

Fluorescent fusion proteins were transiently expressed in *Nicotiana benthamiana* leaf tissue using *Agrobacterium*-mediated infiltration. *Agrobacterium* cells were grown in LB media overnight. Cultures were diluted ten-fold into fresh LB and grown for another 24 hrs. Samples were centrifuged, the pellet was washed once in infiltration media (10 mM MES pH 5.6, 10 mM MgCl_2_, 200 μM acetosyringone) and resuspended to a final OD_600_ of 0.5. Six-week old *N. benthamiana* plants were inoculated with a 1 ml needleless syringe. Leaf discs were viewed using a confocal microscope 1-4 days after inoculation. AtGAAP-GFP/YFP fusion constructs were expressed alone, or co-expressed with organelle markers generated previously (Nelson *et al*., 2007). For colocalisation studies, bacterial suspensions were mixed in a 1:1 ratio before infiltration. Organelle markers used in this study and the corresponding Arabidopsis Biological Resource Centre (ABRC) stock numbers are as follows: Golgi-CFP (CD3-962), Golgi-YFP (CD3-966).

### Confocal microscopy

Leaf discs were mounted on glass slides with water. GFP, YFP, and CFP fluorescence was observed with a Leica DMIRE2 confocal microscope after excitation at 488 nm, 514 nm and 458 nm, respectively. Images were captured and analysed using Leica confocal software (Version 2.61; Leica Microsystems). For colocalization studies sequential scanning alternating between CFP and YFP/GFP channels was performed to avoid crosstalk between the fluorosphores.

### Immunoblot analysis of *Nicotiana benthamiana* tissue

Ivoclar Vivadent Silamat S6 mixer was used to grind frozen leaf discs, followed by protein extraction in 2% SDS, 50 mM Tris-HCl pH 7.5 and protease inhibitors (Sigma P9599, 1:100) at 37°C for 20 minutes. Total protein (50 µg) was run on a 15% SDS polyacrylamide gel, and electroblotted onto a polyvinylidene difluoride (PVDF) membrane. Immunological reactions were performed with anti-GFP mouse monoclonal antisera (1:1000) before detection with a horseradish peroxidase (HRP)-conjugated goat anti-mouse secondary antibody (1:10000).

### Accession numbers

Sequence data can be found in the GenBank/EMBL libraries under the following accession numbers: At4g14730 (*AtGAAP1*), At3g63310 (*AtGAAP2*), At4g02690 (*AtGAAP3*), At4g15470 (*AtGAAP4*), and At1g03070 (*AtGAAP5*).

## RESULTS

### The GAAP gene family is expanded in plants

Both vGAAP and hGAAP belong to the Transmembrane Bax Inhibitor-1 Motif-containing (TMBIM) or Lifeguard (LFG) protein family (Gubser *et al*., 2007; Carrara *et al*., 2012; Saraiva *et al*., 2013a; Saraiva *et al*., 2013b; Carrara *et al*., 2015). Mammals possess six TMBIM subtypes (TMBIM1-6) (Hu *et al*., 2009; Rojas-Rivera and Hetz, 2015). Only two subtypes, GAAP (LFG4/TMBIM4) and BI-1 (TMBIM6), are conserved in plants but several plant species show an intriguing expansion of the GAAP protein family (Gubser *et al*., 2007; Gamboa-Tuz *et al*., 2018). The Arabidopsis, rice (*Oryza sativa*) and the basal angiosperm *Amborella trichopoda* genomes contain five, seven and two putative *GAAP*s, respectively (Figure 1). Arabidopsis *GAAP*s were designated as *AtGAAP1-5* (**Supplementary Table S1**; Guo *et al*., 2018) after the founding members of this gene family (Gubser *et al*., 2007). Arabidopsis GAAP paralogues show 48-82% amino acid sequence identity with each other (**Supplementary Table S2**). Based on phylogenetic analysis AtGAAP2, AtGAAP3 and AtGAAP5 form a distinct clade with high sequence similarity while AtGAAP4 and AtGAAP1 are distinct from both the AtGAAP2/3/5 group and each other (Figure 1**; Supplementary Table S2**). AtGAAP4 shares the highest level of identity with hGAAP (40%), followed by AtGAAP1 (37%), AtGAAP2 (36%), AtGAAP5 and AtGAAP3 (34%; **Supplementary Table S2**). Protein length, transmembrane domain structure and hydrophobicity profiles are also evolutionarily well conserved between Arabidopsis, viral and human GAAPs (**Supplementary Table S1; Supplementary Figures S1A and S1B**; Gubser *et al*., 2007). This includes as little as two to three amino acid difference in protein length between GAAP from camelpox virus (CMLV) and AtGAAP1 and AtGAAP2.

**Figure 1.**
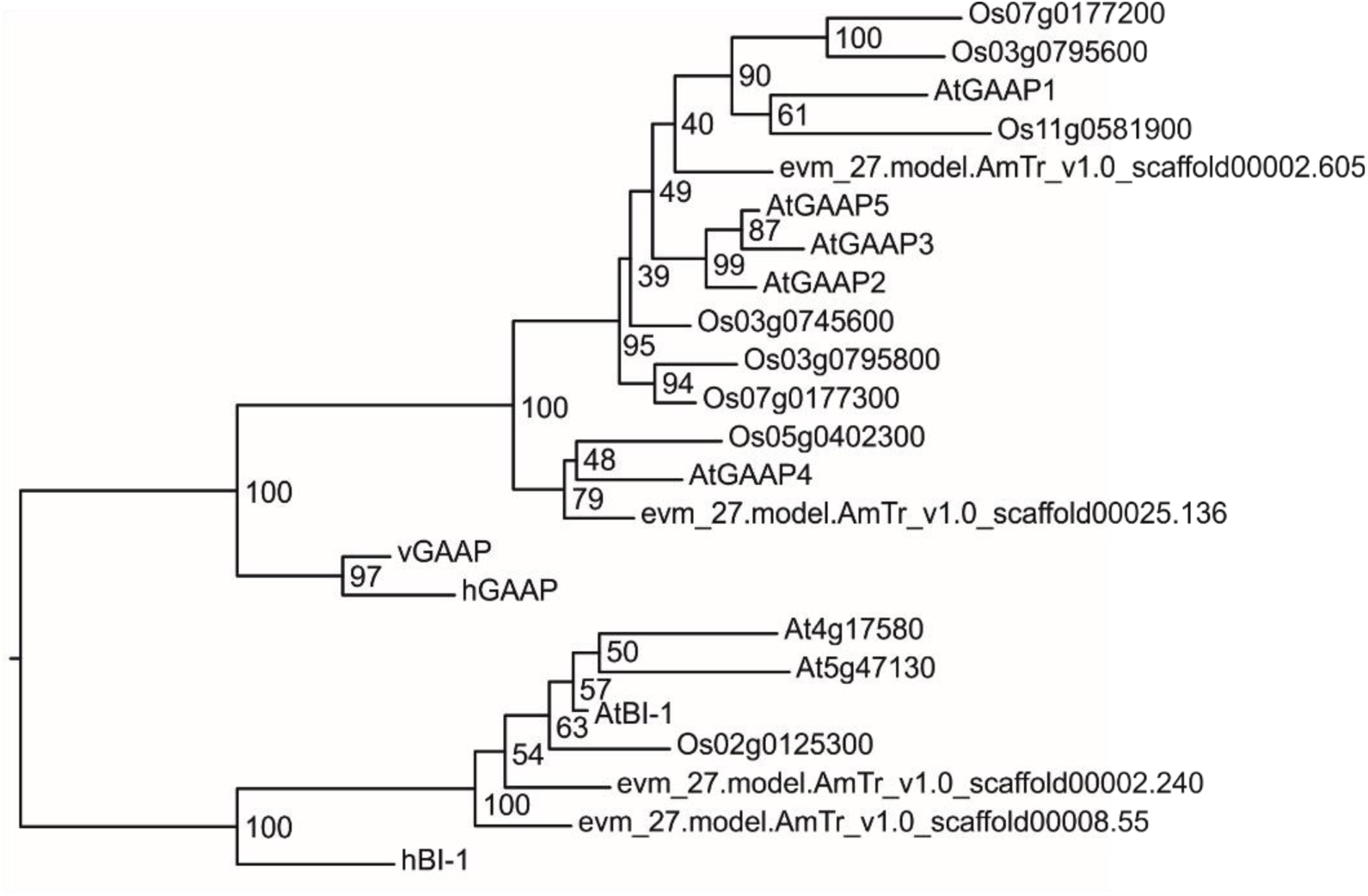
Phylogenetic relationship of GAAP and BI-1 proteins. Maximum-Likelihood tree of GAAP and BI-1 sequences from *Homo sapiens*, camelpox virus, *Arabidopsis thaliana*, *Oryza sativa* and *Amborella trichopoda*. Numbers next to the nodes represent bootstrap values from 1000 replicates. hGAAP, human GAAP; hBI-1, human BI-1; vGAAP, viral GAAP: AtGAAP1-5, Arabidopsis GAAP1-5; AtBI-1, Arabidopsis BI-1; other sequences are labeled according to the gene identifiers.

### Structural models of AtGAAPs define a putative ion channel pore

The structure of a bacterial orthologue of GAAP and BI-1, BsYetJ, has been determined (Chang *et al*., 2014). This structure revealed seven transmembrane domains (TMDs) and a transmembrane pore capable of assuming open and closed conformations (Chang *et al*., 2014). This supports the reported functions of GAAPs and BI-1 as ion channels or antiporters (Kim *et al*., 2008; Ahn *et al*., 2009; Lee *et al*., 2010; Bultynck *et al*., 2014; Carrara *et al*., 2015). Like BsYetJ, AtGAAPs were predicted to contain seven TMDs (**Supplementary Figure S1**). Sequence alignments of AtGAAPs and BsYetJ suggest a similar arrangement of predicted TMDs and substantial (25-32% overall) sequence identity, which was particularly high (37-57%) in the predicted TMD7 region (**Supplementary Figure S1**).

Using the crystal structures of BsYetJ (Chang *et al*., 2014) as templates, we constructed homology models of AtGAAPs in putative closed (Figure 2 and **Supplementary Figures S2-S5**, left columns) and open states (Figure 2 and **Supplementary Figures S2-S5**, middle columns). The models of AtGAAPs contain seven TMDs, with TMD7 at the core of each structure similar to BsYetJ (Chang *et al*., 2014), BI-1 and vGAAP (Carrara *et al*., 2012; Carrara *et al*., 2015). In BsYetJ salt bridges between two aspartates in the pore (D171 and D195), and between D171 and the basic residue R60 in TMD2 stabilize the closed state (Chang *et al*., 2014). Disruption of these salt bridges opens the pore by displacing TMD2 (Chang *et al*., 2014). The AtGAAPs contain aspartate residues corresponding to D171 and D195 of BsYetJ (Figure 2A; **Supplementary Figures S1B; S2A-S5A**), and all AtGAAPs except AtGAAP1 have a positively charged histidine at the position corresponding to R60 of BsYetJ (**Supplementary Figure S1B**). The models suggest that the conserved aspartates are positioned towards the centre of the pore and that the equivalents of BsYetJ R60 are positioned near the cytosolic end of TMD2 (Figure 2 and **Supplementary Figures S2-S5**), similar to their positions in BsYetJ. The closed-state models of AtGAAPs support the possibility that interactions between the aforementioned residues leads to blocking of a transmembrane pore (Figure 2B **and** 2C and **Supplementary Figures S2B-S5B; S2C-S5C**, left and right columns) as in BsYetJ (Chang *et al*., 2014). In the open-state models of AtGAAPs, the residues are spaced further apart (Figure 2B **and** 2C and **Supplementary Figures S2B-S5B; S2C-S5C**, middle columns) consistent with disruption of potential salt bridges. Surface models of AtGAAPs show a pronounced cavity extending along the axis of the pore (Figure 2D **and Supplementary Figure S6**). In AtGAAP2 and AtGAAP3, a continuous pore that fully traverses the membrane is observed in the open state (Figure 2D **and Supplementary Figure S6,** middle and right columns). These results suggest that AtGAAPs may be ion channels or exchangers.

**Figure 2.**
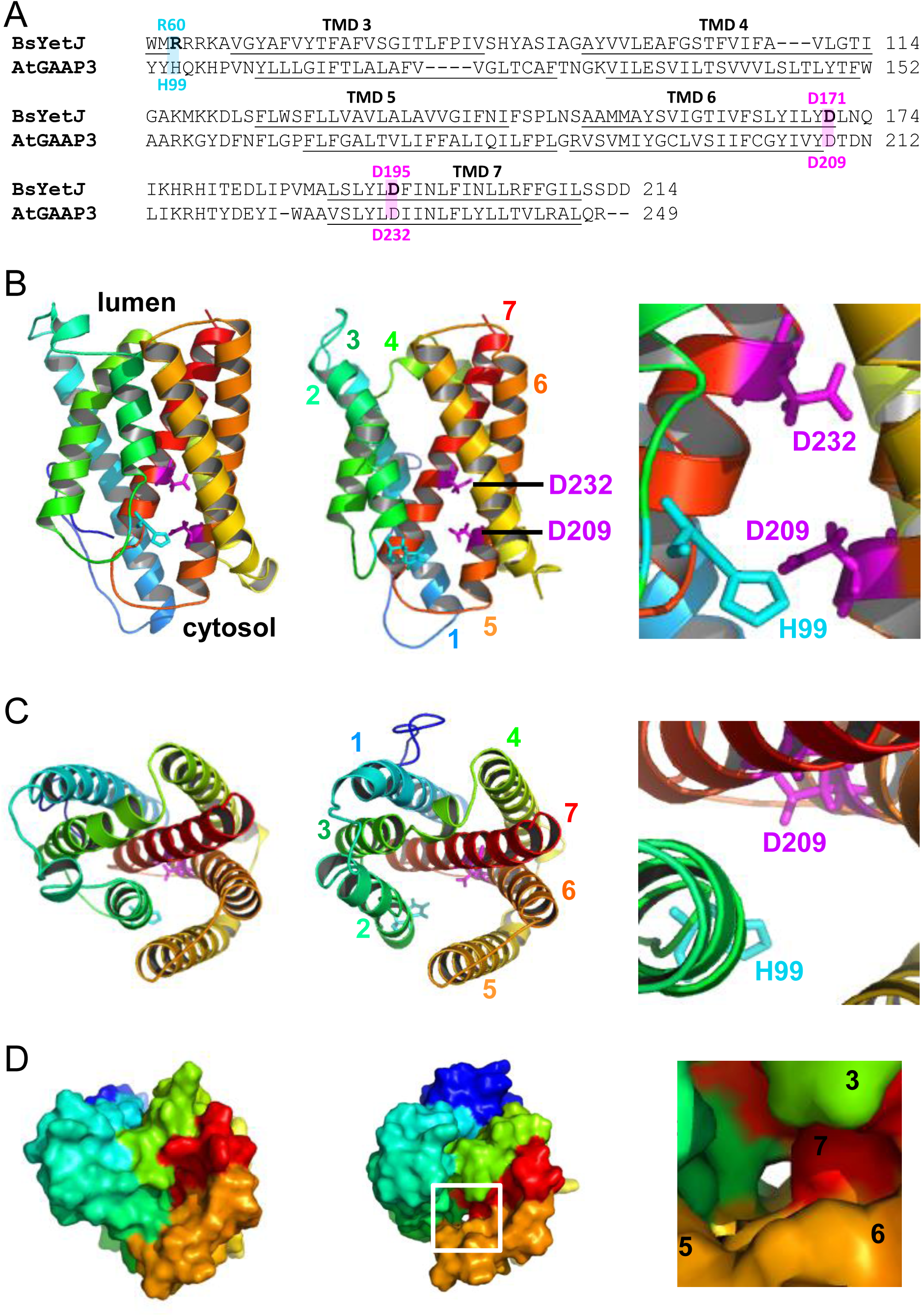
Structural models of AtGAAP3 define a putative ion channel pore. Homology models of AtGAAP3 based on the crystal structures of BsYetJ. **(A)** A sequence alignment of BsYetJ with AtGAAP3. Residues discussed in the text are labeled and predicted transmembrane domains (TMDs) are underlined. **(B-D)** Models of AtGAAP3 in the closed (left column) and open (middle column) states, viewed from the membrane (**B**) or from the lumen of the Golgi apparatus/vacuole **(C-D)**. The side-chains of residues discussed in the text are coloured magenta (equivalents of BsYetJ-D171/D195) and cyan (equivalents of BsYetJ-R60). Helices are coloured for clarity: TMD1 (dark blue), TMD2 (light blue), TMD3 (green), TMD4 (yellow), TMD5 (light orange), TMD6 (dark orange) and TMD7 (red). The insets (right column) show enlarged regions of the closed-state models **(A-B)** or the open-state model **(D)**. Residues discussed in the text are labeled and the TMDs are numbered for clarity.

### AtGAAPs homo-oligomerise and form ion channels in lipid bilayers

Since structural modelling of AtGAAPs predicted the presence of a channel-like pore, we investigated their ability to form ion channels in artificial lipid bilayers. AtGAAP3 was chosen for this analysis because after expression in *Saccharomyces cerevisiae*, detergent extraction and purification it showed a high degree of stability and remained in a non-aggregated state (Figure 3A-C). Recombinant AtGAAP3 was then used for electrophysiological recording after reconstitution into lipid bilayers. Size exclusion chromatography (SEC) (Figure 3A) and non-reducing SDS-PAGE (Figure 3B) of purified AtGAAP3 revealed the expected presence of three or more oligomeric states, similar to vGAAPs and hBI-1 (Saraiva *et al*., 2013b; Carrara *et al*., 2015). Purified AtGAAP3 separated into two oligomeric populations by SEC, with the smallest (monomers) eluting last (Figure 3A). This matches the profiles of the previously characterised vGAAP monomer (Figure 3A, dotted line) and a higher state oligomer (likely a dimer) (Carrara *et al*., 2015). Previous work showed that monomeric vGAAP retained anti-apoptotic activity and reduced the Ca^2+^ content of intracellular stores, while the functional contributions of oligomeric vGAAPs are unknown (Saraiva *et al*., 2013b). Both monomeric and oligomeric populations of purified AtGAAP3 were pooled (Figure 3A, bracket) and used for functional reconstitution into bilayers, in order to optimise opportunities for detecting channel activity.

Incorporation of purified AtGAAP3 in giant unilamellar vesicles (GUVs) into artificial planar bilayers (Figure 3D) gave rise to spontaneous openings of single channels (Figure 3E) similar to those reported previously for vGAAP and BI-1 (Carrara *et al*., 2015). These conductances were not observed in untreated lipid bilayers following the addition of GUVs reconstituted in the absence of protein or with 0.002% lauryldimethylamine N-oxide (LDAO) to mimic any possible detergent-induced artefacts (Figure 3E). These results provide the first direct evidence that AtGAAP3 forms a channel.

**Figure 3.**
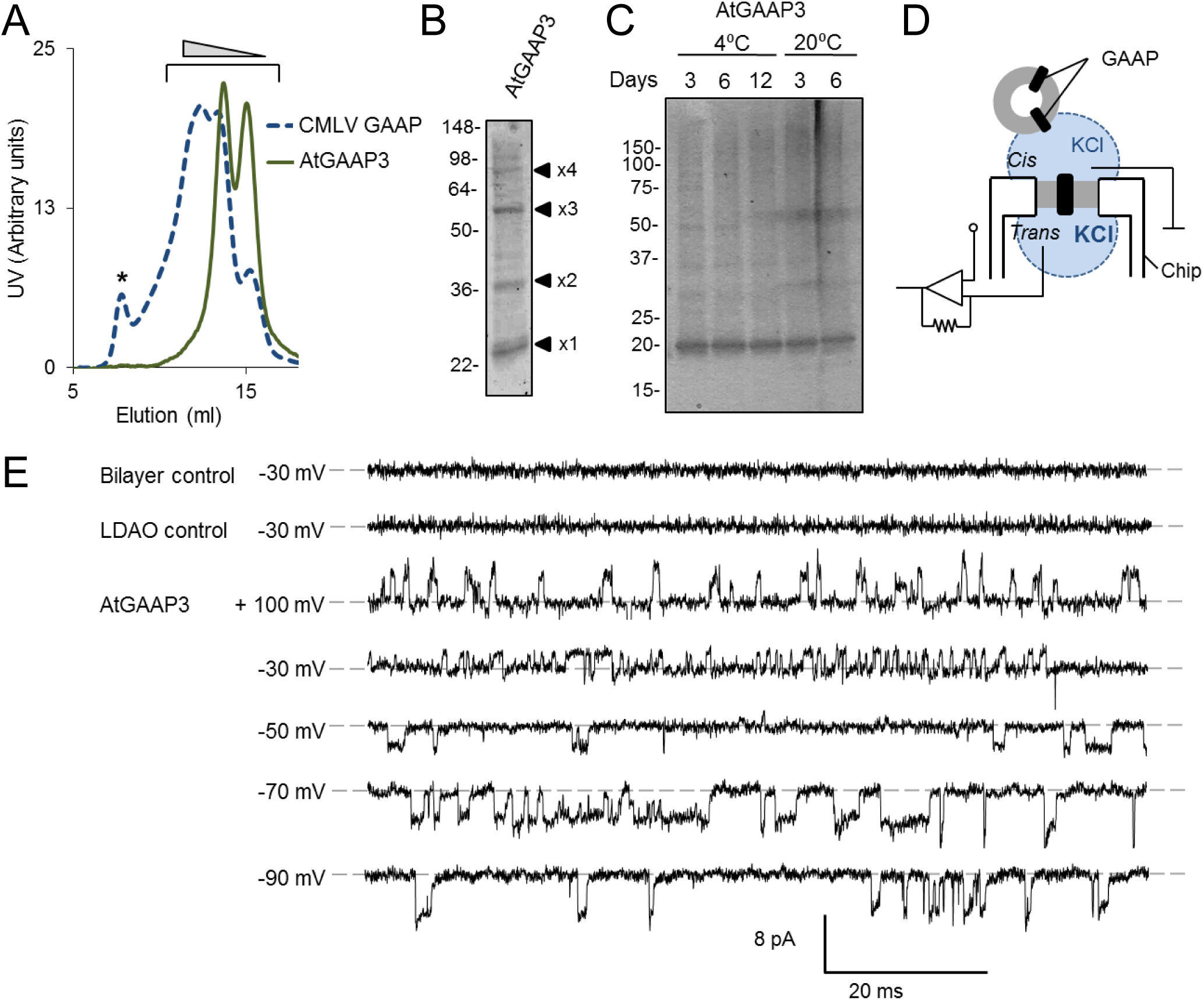
Purified AtGAAP3 exhibits ion channel activity in planar lipid bilayers. **(A-C)** Biochemical analyses of purified AtGAAP3, carried out as described for other GAAPs (Carrara *et al*., 2015). **(A)** Ultraviolet (UV) absorbance profile of purified AtGAAP3 during size exclusion chromatography (SEC) overlapped with previously characterised purified camelpox virus GAAP (CMLV GAAP) (Carrara *et al*. 2015) for comparison. *Corresponds to the protein aggregation peak and the ramp indicates the directional increase in size of eluting GAAP oligomers. Fractions corresponding to monomeric and oligomeric populations of AtGAAP3 (indicated in bracket) were pooled, concentrated, and **(B)** their contents were analysed by non-reducing SDS-PAGE and Coomassie staining. The expected positions of the monomeric (x1) and oligomeric proteins (x2, x3 and x4) are shown. **(C)** The stability and aggregation of the purified AtGAAP3 was assessed over a period of 12 days at 4°C and 20°C and proteins were visualised by Coomassie stain. **(D-E)** Electophysiological analyses of AtGAAP function, were carried out as described previously for other GAAPs (Carrara *et al*., 2015). (**D**) Conditions of the bilayer chamber used. A planar lipid bilayer is formed across a µm-sized aperture within the chip. Purified protein is incorporated into the bilayer by adding protein reconstituted GUVs to the cis chamber (ground). The KCl concentration is greater in the *trans* chamber relative to the *cis* chamber. **(E)** Electrophysiological recordings from artificial lipid bilayers reconstituted with purified AtGAAP3 show spontaneous channel openings. Representative current traces were recorded at the indicated holding potentials, which are expressed as the potential on the *cis* side relative to the *trans* side (n = 2 independent experiments). Downward deflections represent positive ions flowing from the *trans* to the *cis* side of the bilayer. The lipid bilayer alone (n = 35) or after addition of GUVs reconstituted in the presence of LDAO (n = 10) were used as negative controls. The dotted line indicates the closed state.

### AtGAAPs rescue yeast from apoptosis

Since hGAAP, vGAAP and BI-1 inhibit apoptosis (Xu and Reed, 1998; Gubser *et al*., 2007), we tested whether AtGAAPs also had anti-apoptotic activity. A strain of *S. cerevisiae* (termed Bax_2_) was generated that expressed the pro-apoptotic protein Bax (Oltvai *et al*., 1993) under the control of a galactose-inducible promoter. This Bax_2_ strain showed a dramatic reduction of growth on galactose-containing medium compared to a control strain (GFP_2_) (Figure 4), consistent with an induction of apoptosis in the Bax_2_ strain. Expression of AtGAAPs or AtBI-1 in the Bax_2_ strain markedly enhanced growth after induction of Bax expression, relative to the control strain expressing GFP (Figure 4). The enhanced growth was greater for AtGAAP1-4 than for AtGAAP5, the latter only showing significantly enhanced growth after longer periods (10 days) of growth (Figure 4). These results demonstrate that AtGAAPs exert an anti-apoptotic effect in yeast, similar to BI-1 (Xu and Reed, 1998).

**Figure 4.**
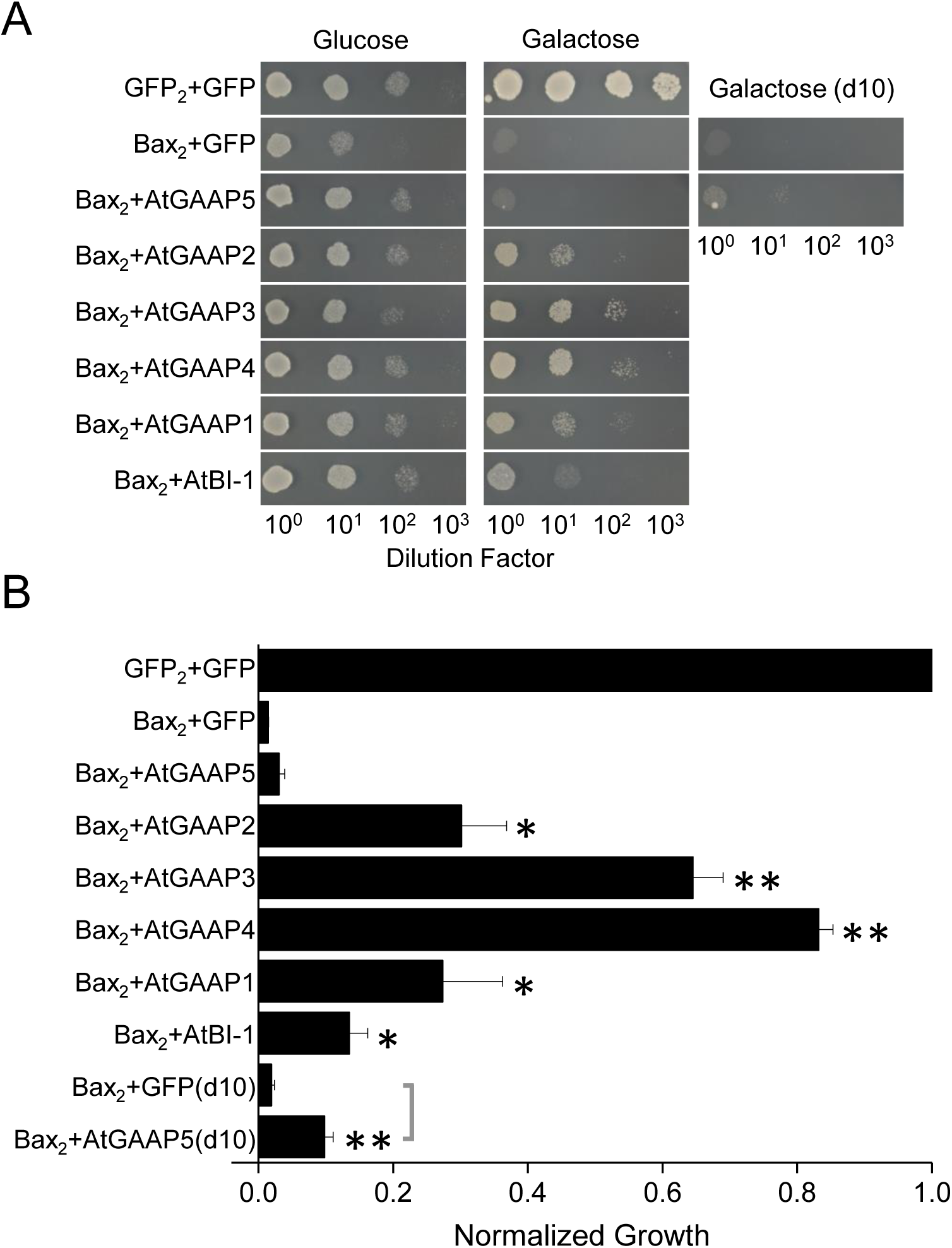
AtGAAPs rescue yeast from Bax-mediated apoptosis. (**A**) Representative growth is shown after 2 days (glucose) or 5 days (galactose), and the inset shows growth after 10 days (galactose) for the slowest growing transformants. (**B**) Mean data from independent transformations (n=3), showing normalized growth on SDgalactose-His-Leu-Ura. Data are represented as mean ± SEM and statistical significance is indicated at \**p*<0.05 and \*\**p*<0.01 levels, using one-way ANOVA and Dunnett’s multiple comparison test.

### AtGAAPs localize mainly to the Golgi *in planta*

Human and viral GAAPs localize mainly to the Golgi (Gubser *et al*., 2007). To study the subcellular localization of Arabidopsis GAAPs *in planta*, AtGAAPs tagged with fluorescent proteins (GFP or YFP) were transiently expressed in *Nicotiana benthamiana* leaves and analysed by confocal microscopy. Expression of the fluorescently-tagged AtGAAPs in the Bax_2_ yeast strain confirmed that they retained their anti-apoptotic effects (**Supplementary Figure S7**), consistent with previous studies using tagged BI-1 (Xu and Reed, 1998) and GAAPs (Gubser *et al*., 2007; de Mattia *et al*., 2009; Saraiva *et al*., 2013a; Saraiva *et al*., 2013b; Carrara *et al*., 2015). At the modest expression levels achieved within one to two days after transformation of *N. benthamiana*, fluorescence localized to small, punctate structures distributed throughout epidermal cells, and colocalized with a Golgi-CFP marker (Figure 5A) (Nelson *et al*., 2007).

**Figure 5.**
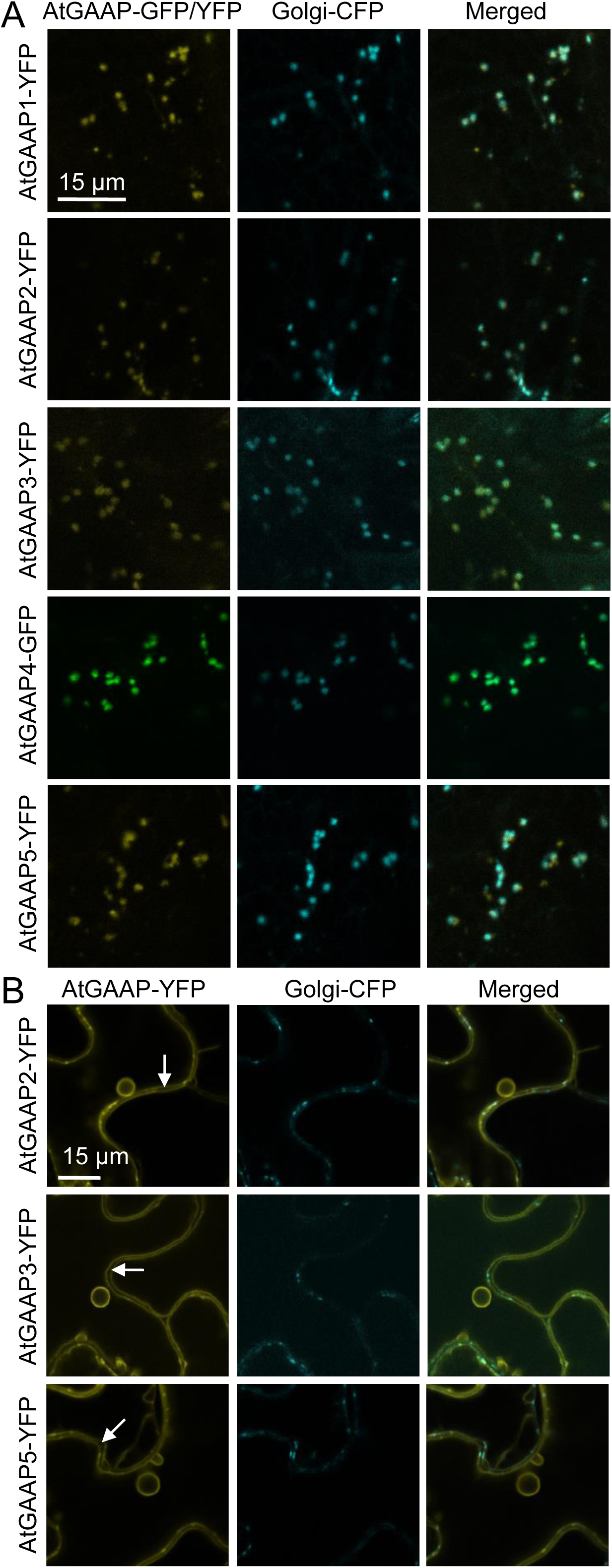
Subcellular localisation of AtGAAP1-5:GFP/YFP fusion proteins in *Nicotiana benthamiana* leaf epidermal cells. Leaves were co-infiltrated with *Agrobacterium* carrying AtGAAP1-5:GFP/YFP and Golgi-CFP marker constructs and imaged using a confocal laser scanning microscope two days **(A)** and three days **(B)** after inoculation. YFP/GFP (left column) and CFP (middle column) signals and merged images (right column) are shown. Scale bars are as indicated. Arrows denote cell wall space between two neighbouring cells. Experiment was repeated at least three times and representative images are shown.

At high expression levels, human and viral GAAPs are also expressed in the ER (Gubser *et al*., 2007). At the higher expression levels achieved three to four days after transformation, AtGAAP1-YFP and AtGAAP4-GFP colocalised exclusively with the Golgi-CFP marker. However, the distributions of YFP-tagged AtGAAP2/3/5 were similar to that of tonoplast proteins (Saito *et al*., 2002; Nelson *et al*., 2007), with a clearly visible cell wall space between two neighbouring cells (Figure 5B, arrows) and ring-like structures (Saito *et al*., 2002). Some colocalisation with the Golgi stacks was also maintained at these later time points for YFP-tagged AtGAAP2/3/5 (Figure 5B). We conclude that like hGAAP and vGAAP, AtGAAPs are expressed primarily in the Golgi at low levels of expression. At higher expression levels, AtGAAP1 and AtGAAP4 remain within the Golgi, whereas AtGAAP2/3/5 are expressed mainly at the tonoplast.

### *AtGAAP*s show differential spatial expression and abundance of transcript

To gain further insight into the possible functions of the *AtGAAP* family, we fused the *AtGAAP* promoters (*proAtGAAP*) to the β-glucuronidase (GUS) reporter gene (*uidA*) and generated transgenic lines. Distinct expression patterns for *AtGAAP1-5* could be observed both spatially and in transcript abundance based on the pattern and intensity of GUS staining between the transgenic lines (Figure 6A**; Supplementary Figure S8**). *AtGAAP4* showed the strongest and most uniform expression throughout the inflorescence and leaf tissues. *AtGAAP2* expression was also comparatively high in both tissue types while *AtGAAP1* and *AtGAAP5* expression was markedly lower. No GUS staining was observed in plants containing the *proAtGAAP3::uidA* transgene (Figure 6A**; Supplementary Figure S8)**.

**Figure 6.**
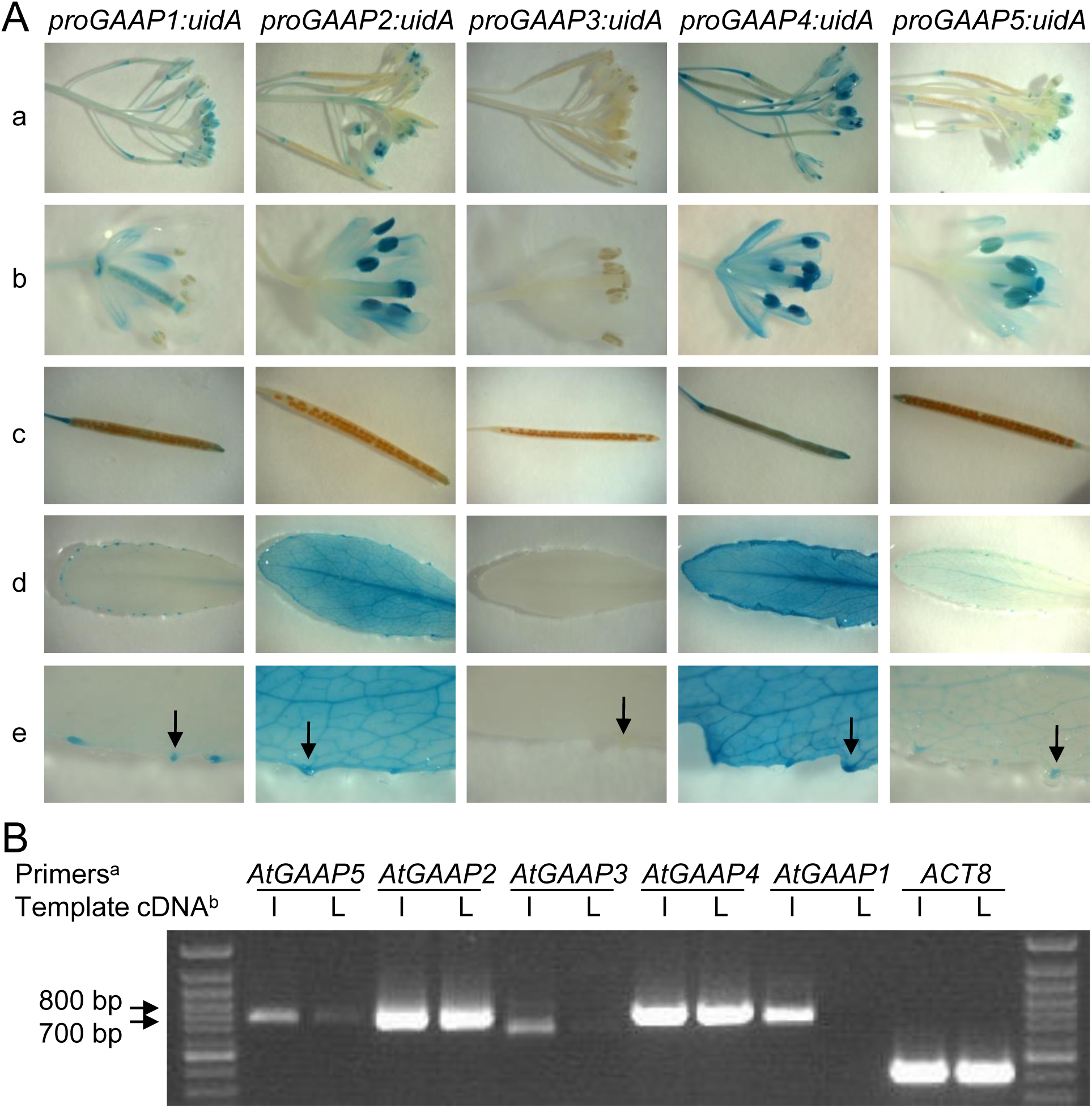
*AtGAAP* gene expression analysis. **(A)** Histochemical analysis of GUS expression in *proAtGAAP1-5::uidA* transgenic plants. Tissue of six-week old Arabidopsis plants expressing *AtGAAP* promoter-GUS fusions was subjected to histochemical staining for GUS activity. 1800 bp, 1800 bp, 1694 bp, 1794 bp and 1803 bp of promoter region was used for *AtGAAP1* to *AtGAAP5*, respectively. **(a)** Inflorescence tissue containing flowers and young siliques, **(b)** individual flower, **(c)** silique, **(d)** rosette leaf, **(e)** magnification of rosette leaf showing staining in veins and hydathodes. Hydathodes are indicated with arrowheads. The expression of *uidA* is driven by *AtGAAP* promoters as indicated. 12 transgenic lines were analysed for each construct and images of representative lines are shown. **(B)** RT-PCR analysis of *AtGAAP* expression. RT-PCR analysis of *AtGAAP1*, *AtGAAP2*, *AtGAAP3*, *AtGAAP4* and *AtGAAP5* expression in five-week old wild type Col-0 inflorescence and leaf tissue. *ACTIN8* (*ACT8)* expression provided a control for RT-PCR. ^a^ Primer pairs used were specific for a particular *AtGAAP* paralogue, or *ACT8* as indicated. ^b^ Total RNA was extracted from either inflorescence (I) or rosette leaf (L) tissue of wild type Col-0 plants as indicated.

*AtGAAP4* was expressed widely in the reproductive tissues, including the pistil, stamen, sepals, petals, siliques and inflorescence stems (Figure 6A**; Supplementary Figure S8)**. *AtGAAP2* and *AtGAAP5* expression was elevated particularly in the anther, including the pollen grains, and the tip of the style and stigma. Unlike the other *AtGAAP* paralogues, *AtGAAP1* was expressed distinctly in the ovules within the pistil and also highly expressed in the petal and sepal abscission zones. In the vegetative tissues, *AtGAAP2* and *AtGAAP4* were highly expressed throughout the rosette with particularly intense GUS staining observed in the leaf vasculature and the hydathodes (Figure 6A). Similar spatial pattern of expression was observed for *AtGAAP5* although at much reduced amounts while *AtGAAP1* showed hydathode-specific expression in the leaves.

To complement the histological analysis of *AtGAAP* promoter activity by GUS staining, the abundance of *AtGAAP* transcripts in leaf and inflorescence tissues was analyzed using semi-quantitative RT-PCR. *AtGAAP2* and *AtGAAP4* showed high transcript abundance in leaves and flowers (Figure 6B). Transcript levels of *AtGAAP1*, *AtGAAP3* and *AtGAAP5* were substantially lower compared to *AtGAAP2* and *AtGAAP4.* The abundance of *AtGAAP1, 3* and *5* transcripts was higher in inflorescence tissue compared to leaves (Figure 6B). Analysis of publicly available expression data in Genevestigator (Hruz *et al*., 2008; Zimmermann *et al*., 2004) supported the results from both *ProAtGAAP*::uidA and RT-PCR analyses (**Supplementary Fig. S9**). We conclude that *AtGAAP*s show widespread expression in Arabidopsis, and that relative transcript abundance and spatial expression pattern in plant tissues differs between the *AtGAAP* paralogues.

### *atgaap* single, double and triple mutants show no obvious growth defects

Gene expression analysis indicated that all *AtGAAPs* are expressed in Arabidopsis. In order to investigate the physiological functions of individual AtGAAPs we isolated homozygous T-DNA insertion mutants in the Colombia-0 (Col-0) background for all *AtGAAP*s. Absence of *AtGAAP1*, *AtGAAP3*, *AtGAAP4* and *AtGAAP5* transcripts suggested that the plants were loss-of-function mutants (*atgaap1* - SALK_46652, *atgaap3* - GABI186E10, *atgaap4* - SAIL 151_F11, *atgaap5* - SALK_066103; **Supplementary Figure S10A-B**). The only available T-DNA line for *AtGAAP2* in the Col-0 background, SALK_52507, showed *AtGAAP2* expression levels similar to wild-type (**Supplementary Figure S10A-B**), but no *AtGAAP2* transcript was detected in a transposon-tagged line in the Landsberg *erecta* (L*er*) background, *atgaap2* - GT_93791. Homozygous mutant plants showed no obvious growth phenotypes and were indistinguishable from wild-type plants when grown under either 10- or 16-hour day regimes (**Supplementary Figure S11**). The mutants were fertile and produced seeds that germinated and developed normally.

Since the lack of phenotypes in *atgaap* single mutants could be the result of genetic redundancy between AtGAAP subtypes, *atgaap* double and triple mutants were generated. We were able to isolate all *atgaap* double mutant combinations except *atgaap1atgaap4*. The short genetic distance between *AtGAAP1* and *AtGAAP4* is the likely reason for not obtaining a double mutant for these genes. Furthermore, we generated *atgaap2atgaap3atgaap4, atgaap2atgaap3atgaap5*, and *atgaap2atgaap4atgaap5*. Intriguingly, no obvious developmental defects were observed in any of the double or triple mutants (**Supplementary Figure S11**). As expected, *atgaap2* double and triple mutants, with mixed Col-0 and L*er* background, displayed varying characteristics of parental ecotypes, similarly to offspring of crosses between the wild-type plants. We conclude from our results that AtGAAP subtypes may display a high degree of functional redundancy in Arabidopsis under normal growth conditions, or that AtGAAPs regulate responses to specific environmental stimuli or stresses.

### Over-expression of AtGAAP1-YFP induces cell death of *Nicotiana benthamiana* leaves and severe growth defects in Arabidopsis

As loss-of-function mutants did not provide insights into AtGAAP function, we studied the effects of transient and stable over-expression of AtGAAPs in *N. benthamiana* and Arabidopsis, respectively. Transient expression in *N. benthamiana* leaf tissue has been employed successfully to study the role of proteins in the induction and inhibition of cell death (Yoshioka *et al*., 2006; Urquhart *et al*., 2007; Yang *et al*., 2007; Baxter *et al*., 2008; Chin *et al*., 2010; Xu *et al*., 2017). Transient over-expression of AtGAAP1-YFP in *N. benthamiana* led to visible symptoms of cell death 2-3 days after inoculation (Figure 7A-B). Conversely, over-expression of fluorescently-tagged AtGAAP2-5, Golgi-YFP marker (Nelson *et al*., 2007) or the fluorescent tags alone did not induce cell death (Figure 7A-B). Immunoblot analysis confirmed transgene expression with YFP-tagged AtGAAP2/3/5, the Golgi-YFP marker and YFP showing the greatest abundance and AtGAAP1-YFP present at lower levels (Figure 7C).

**Figure 7.**
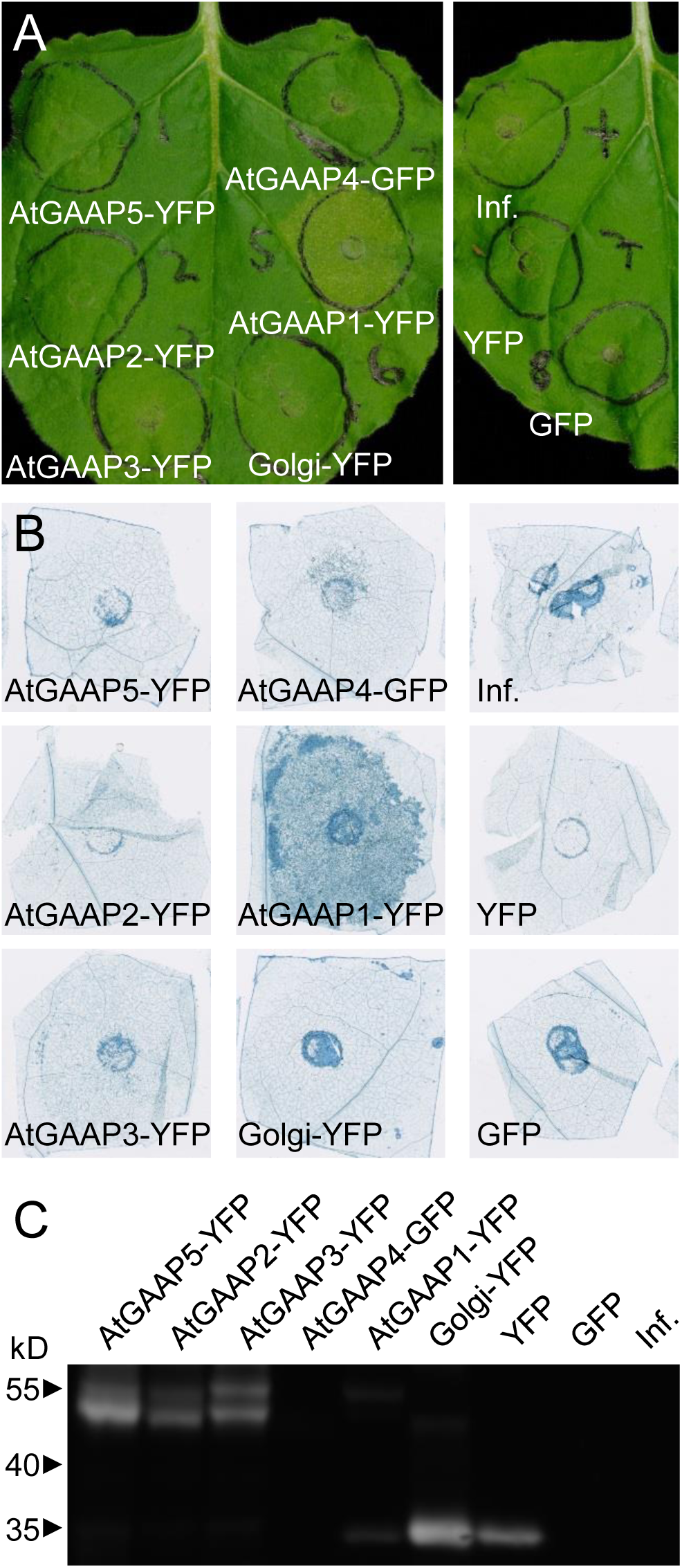
Over-expression of AtGAAP1-YFP induces cell death upon transient expression in *Nicotiana benthamiana* leaf tissue. **(A)** Leaf areas marked with circles were infiltrated with Agrobacterium carrying AtGAAP1/2/3/5-YFP, AtGAAP4-GFP, Golgi-YFP marker, YFP and GFP constructs or infiltration media (Inf.) as indicated. Shown are representative leaves at three days post inoculation (dpi). **(B)** Inoculated leaf areas were excised from the leaves and subjected to trypan blue staining for visualization of cell death at three dpi. **(C)** Immunoblots showing expression of fusion proteins in inoculated leaf areas at three dpi. Experiments were repeated at least three times, representative images are shown.

In order to address whether the induction of cell death by AtGAAP1 was also apparent in Arabidopsis, transgenic plants stably expressing AtGAAP1-YFP driven by the strong constitutive promoter 35S in the *atgaap1* mutant background were generated. Plant morphology was dramatically altered in the plants over-expressing AtGAAP1-YFP (Figure 8). Rosettes were small in size compared to *atgaap1* plants, and over-expression plants displaying a range of sizes could be identified. A representative panel of 5-week old plants displaying varying degrees of the developmental phenotype is shown in Figure 8A. Lower rosette leaves of plants over-expressing AtGAAP1-YFP exhibited early senescence compared to *atgaap1* plants (Figure 8B-C). In plants over-expressing AtGAAP1-YFP, the oldest rosette leaves that lacked visible signs of yellowing, as indicated by arrows in Figure 8C, had brown lesions (Figure 8D). These lesions were clearly identified as regions of dead cells with trypan blue staining (Figure 8E). While trypan blue staining for dead cells was most pronounced around the site of the lesions, staining was also observed throughout these leaves. No cell death was observed in *atgaap1* leaves of similar age (Figure 8F). The distinct punctate pattern of YFP fluorescence typical of the Golgi was observed in plants over-expressing AtGAAP1-YFP, indicating that the observed phenotypes were likely due to accumulation of the recombinant protein in its native location (**Supplementary Figure S12**).

**Figure 8.**
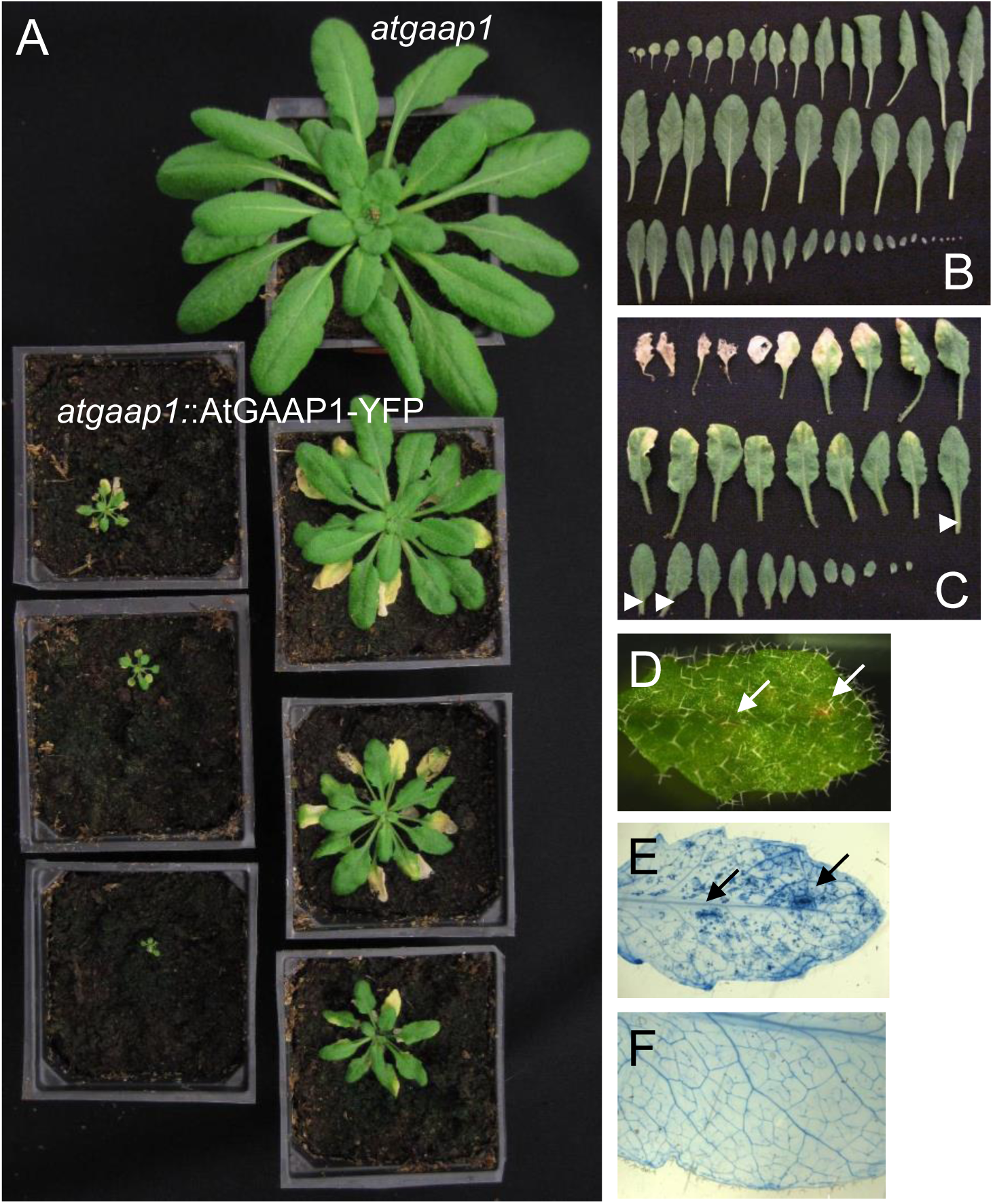
Over-expression of AtGAAP1-YFP leads to dwarfism, enhanced senescence and development of spontaneous lesions on the rosette leaves in Arabidopsis. **(A)** An *atgaap1* plant and a panel of *atgaap1* plants expressing AtGAAP1-YFP (*atgaap1*::AtGAAP1-YFP) as indicated photographed at five weeks. Expression of AtGAAP1-YFP led to a dwarf phenotype and induced senescence of lower rosette leaves. Two to five transgenic lines deriving from three independent transformations were analysed. All displayed similar phenotypes. **(B-C)** Rosette leaves of a five-week old *atgaap1* plant **(B)** and an *atgaap1* plant expressing AtGAAP1-YFP **(C)**. **(D)** The oldest rosette leaves of plants expressing AtGAAP1-YFP without signs of yellowing (indicated by arrowheads in panel C) developed brown lesions (arrows) **(E)** The leaf in **(D)** after trypan blue staining for cell death. Brown lesions in **(D)** are clearly visible as regions of dead cells in **(E)**. **(F)** Trypan blue stained leaf of an *atgaap1* plant. No cell death was visible. Leaves of the same age were used for **(D-F).**

In summary, these data show that ectopic over-expression of AtGAAP1-YFP, but not AtGAAP2-5, induces cell death in *N. benthamiana* leaf tissue. Over-expression of AtGAAP1-YFP in Arabidopsis leads to dwarfism, early senescence and ectopic cell death. This supports a role for plant GAAPs in the control of cell death *in planta*.

## DISCUSSION

### Arabidopsis GAAPs are channel proteins that localize to the Golgi and tonoplast

The homology models based on the structure of BsYetJ suggested that AtGAAPs form transmembrane pores that function as ion channels or exchangers. This is similar to previous proposals that BI-1 (Bultynck *et al*., 2014), vGAAPs (Carrara *et al*., 2015) and BsYetJ (Chang *et al*., 2014) form ion channels or exchangers that mediate Ca^2+^ flux across membranes. The aspartate pair (Asp171-Asp195) in BsYetJ is involved in hydrogen bonding responsible for the reversible, pH-dependent transitions between open and closed conformation of BsYetJ that are associated with Ca^2+^ leak across the membrane (Chang *et al*., 2014). Importantly, the conservation of this di-aspartyl pH sensor in AtGAAPs as well as viruses and vertebrates (Carrara *et al*., 2015) suggests a similar regulatory mechanism for GAAPs from bacteria, animals, viruses and plants. Mutagenesis, in conjunction with functional assays, will be required to verify the importance of the residues concerned.

vGAAP and hBI-1 form ion channels and vGAAP is selective for cations (Carrara *et al*., 2015). In this study, we demonstrate that purified AtGAAP3 exhibits ion channel activity in planar lipid bilayers. To our knowledge this is the first direct evidence that a plant TMBIM family member exhibits ion channel functions. The relative selectivity of the AtGAAPs for cations remains to be determined. Although it has been suggested that plant BI-1 may be directly involved in Ca^2+^ transport across the ER membrane (Ishikawa *et al*., 2011), experimental evidence is still lacking. In contrast, Ca^2+^ leak properties reported for the C terminus of BI-1 were conserved in animal but not plant orthologues, suggesting that the Ca^2+^ channel-like activity of BI-1 may have been acquired specifically in the animal lineage (Bultynck *et al*., 2012). Residues that are important for ion conductance (Asp219 and Glu207 of vGAAP) and ionic selectivity (Glu207) of vGAAP (Carrara *et al*., 2015) are conserved in AtGAAPs (**Supplementary Figure S1**), further implying functional conservation.

The possible role of AtGAAPs as a novel class of Ca^2+^-conducting channels in plants is interesting in light of the evidence showing that plants generally show a loss of diversity in the mechanisms for Ca^2+^ influx from the extracellular space and intracellular stores into the cytoplasm compared to animals (Wheeler and Brownlee, 2008; Verret *et al*., 2010; Edel and Kudla, 2015; Marchadier *et al*., 2016; Edel *et al*., 2017); many of the classic animal Ca^2+^ channels like the inositol 1,4,5-trisphosphate receptors (IP_3_Rs) are not present in plant genomes (Wheeler and Brownlee, 2008; Verret *et al*., 2010; Edel and Kudla, 2015). Thus far, only five families of Ca^2+^-permeable influx channels have been shown to function in plants: cyclic nucleotide-gated channels (CNGCs)(Zelman *et al*., 2012), glutamate-like receptors (GLRs)(Price *et al*., 2012), two-pore channels (TPCs) (Morgan and Galione, 2014), mechanosensitive channels (MCAs) (Kurusu *et al*., 2013) and reduced hyperosmolality-induced Ca^2+^ increase channels (OSCAs) (Yuan *et al*., 2014). Reduced diversity of mechanisms for Ca^2+^ influx in plants compared to animals is associated with amplification of specific mechanisms, as many of these gene families show expansion in plants (Edel *et al*., 2017) as is also demonstrated in the case of plant GAAPs (Figure 1).

Like hGAAP and vGAAP (Gubser *et al*., 2007), AtGAAPs localize to Golgi membranes, suggesting a conserved function as Golgi-localized ion channels. The role of the Golgi as an important Ca^2+^ store that contributes to Ca^2+^ signaling in animal cells is established (Pizzo *et al*., 2011). Although information on Ca^2+^ handling by the plant Golgi is scarce (Costa *et al*., 2018), it has been shown that free Ca^2+^ concentration is higher in the Golgi than in the cytosol of plant cells, and abiotic cues can affect luminal Ca^2+^ dynamics (Ordenes *et al*., 2012). The nature and identity of Ca^2+^ channels and transporters of the plant Golgi awaits clarification, although a role for P2A-type ATPase AtECA3 in the transport of Ca^2+^ and Mn^2+^ into the Golgi has been proposed (Mills *et al*., 2008). AtGAAP2/3/5, but not AtGAAP1/4, also localized in the tonoplast (Figure 5). The vacuole is the main storage compartment of Ca^2+^ in plant cells (Peiter, 2011; Costa *et al*., 2018) and many tonoplast Ca^2+^ transporters have been identified (Martinoia *et al*., 2012; Neuhaus and Trentmann, 2014; Costa *et al*., 2018). Ca^2+^ release from the vacuole likely contributes to signaling in plants and here evidence for a role for TPC1 as a tonoplast channel critical for many physiological processes is accumulating (Peiter *et al*., 2005; Carpaneto and Gradogna, 2018). Localization of AtGAAPs to these organelles makes them ideally placed to act as functional Ca^2+^ release channels.

### Arabidopsis GAAPs regulate cell death in yeast cells and *in planta*

Similar to hGAAP and vGAAP (Gubser *et al*., 2007), all five AtGAAPs suppressed cell death induced by Bax-expression in yeast (Figure 4 **and Supplementary Figure S7**). This suggests that the anti-apoptotic effects of GAAPs are conserved between plants, viruses and vertebrates. Various studies in mammals indicate that other members of the TMBIM family also display anti-apoptotic activity (Rojas-Rivera and Hetz, 2015). BI-1 suppresses Bax-induced cell death in yeast (Xu and Reed, 1998), and Bax-induced PCD is suppressed by BI-1 proteins in Arabidopsis, rice, rapeseed and tobacco (Kawai *et al*., 1999; Sanchez *et al*., 2000; Kawai-Yamada *et al*., 2001; Bolduc *et al*., 2003). This conservation of function within the TMBIM family suggests that these proteins target one of the core molecular mechanisms of PCD that is conserved across kingdoms.

Expression of AtGAAP1-YFP in yeast inhibited Bax-induced cell death, whereas expression of this fusion protein in *N. benthamiana* and Arabidopsis induced cell death, demonstrating that AtGAAP1 can function as a positive or negative regulator of cell death depending on the context. Opposite effects of BI-1 in regulating cell death have also been demonstrated. Mammalian and plant BI-1 have been mainly associated with cytoprotective functions (Xu and Reed, 1998; Kawai *et al*., 1999; Sanchez *et al*., 2000; Kawai-Yamada *et al*., 2001; Bolduc *et al*., 2003; Robinson *et al*., 2011). However, AtBI-1 has been reported to induce cell death in mammalian cells (Yu *et al*., 2002) and upon over-expression in *N. benthamiana* (Xu *et al*., 2017). The former effect was speculated to be due to AtBI-1 functioning as a dominant inhibitor of the endogenous mammalian BI-1, while the latter was shown to be dependent on certain autophagy-related proteins. A cell death-promoting phenotype for mammalian BI-1 at low pH has also been reported (Kim *et al*., 2008; Lee *et al*., 2011); over-expression of BI-1 leads to increased Ca^2+^ release from the ER and promotion of cell death in acidic conditions (Kim *et al*., 2008), while BI-1 knock-down had the opposite effects (Lee *et al*., 2011). These studies and the results reported here demonstrate dual and contrasting roles for TMBIM proteins in regulating cell death, depending on the context. The molecular mechanisms underlying these observed effects remain to be characterized in detail but may be related to the function of these proteins as cation or Ca^2+^ channels.

Stable over-expression of AtGAAP1-YFP fusion protein in Arabidopsis led to a severe dwarf phenotype, enhanced leaf senescence and development of spontaneous lesions in leaves (Figure 8). This phenotype resembles that of lesion mimic mutants (LMM), which display spontaneous PCD and have been widely used as models for unraveling cell death signaling pathways (Lorrain *et al*., 2003; Bruggeman *et al*., 2015). Some previously characterized LMMs contain mutations in genes encoding ion channels. *Constitutive expresser of PR genes 22* (*cpr22*) encodes the chimeric cyclic nucleotide-gated ion channel CNGC11/CNGC12, which is a constitutively active form of the putative Ca^2+^ channels responsible for the lesions (Yoshioka *et al*., 2001; Yoshioka *et al*., 2006; Urquhart *et al*., 2007; Chin *et al*., 2010). In contrast, mutations disrupting CNGC2 and CNGC4 in *defence no death1* (*dnd1*) and *dnd2*, respectively, lead to LMM phenotypes (Yu *et al*., 1998; Clough *et al*., 2000; Balague *et al*., 2003; Jurkowski *et al*., 2004; Ahn, 2007; Ali *et al*., 2007). Given the critical role of ion fluxes in the regulation of PCD, and the LMM phenotypes previously recorded for plant ion channel mutants, it is tempting to speculate that the LMM phenotype induced by AtGAAP1-YFP over-expression could be due to unbalanced ionic homeostasis caused by the over-expression of a member of this novel family of ion conducting channels.

### Further physiological roles of AtGAAPs

Our gene expression analysis provided clear evidence for expression of all five *AtGAAP* paralogues. *AtGAAP2* and *AtGAAP4* showed the highest and most uniform abundance of transcript throughout plant tissues. Expression of *AtGAAP1* and *AtGAAP5* was much lower, with increased expression in the inflorescence tissue compared to rosettes. *AtGAAP3* was the least abundant *AtGAAP* isoform according to all methods used. It is noteworthy that in the sexual organs, *AtGAAP2*, *AtGAAP4* and *AtGAAP5* showed prominent expression in the pollen whereas *AtGAAP1* expression was most prominently elevated in the ovary. Taken together, these data suggest that AtGAAPs may fulfil subtype-specific roles in different tissues. The tissue expression analysis will facilitate predictions of possible redundancy between individual AtGAAP subtypes and the processes that they regulate. For example, the abundance of *AtGAAP2*, *AtGAAP4* and *AtGAAP5* transcripts in pollen could indicate the involvement of these proteins in the tapetal PCD that occurs during pollen development (Kurusu and Kuchitsu, 2017), a process in which AtBI-1 may play an inhibitory role (Kawanabe *et al*., 2006).

In order to obtain further insights into the *in vivo* functions of AtGAAPs we analyzed single, double and triple mutants. The lack of developmental phenotypes in the mutants suggested that AtGAAPs may display a high degree of functional redundancy in Arabidopsis, or that AtGAAPs regulate responses to specific environmental stimuli or stresses. Recently, a redundant role for AtGAAP1 to AtGAAP3 in the unfolded protein response and the onset of cell death in response to ER stress was reported (Wang *et al*., 2019; Guo *et al*., 2018). A role for barley GAAP in plant-fungal interaction has also been demonstrated (Weis *et al*., 2013) and this function is conserved in barley BI-1 (Huckelhoven *et al*., 2003; Eichmann *et al*., 2006; Babaeizad *et al*., 2009). Knockdown of AtGAAP1 (AtLFG1) or AtGAAP2 (AtLFG2) also delayed fungal development on Arabidopsis (Weis *et al*., 2013). Intriguingly, AtGAAP2 has also been linked to brassinosteroid signaling (Yamagami *et al*., 2009). Further studies are required to uncover the molecular mechanisms underlying the reported phenotypes and additional phenotypes.

In summary we describe and characterize a family of five GAAPs in Arabidopsis. We demonstrate that they form ion channels and share an evolutionarily conserved function in cell death regulation with GAAPs from animals and viruses. We propose that the physiological role of AtGAAPs in cell death regulation is based on the ability of these Golgi-and tonoplast-localized ion channels to modify the finely balanced ionic homeostasis within the cell. Differential expression of AtGAAP subtypes in different plant tissues suggests that this family of proteins may play roles in a variety of physiological processes.

## SUPPLEMENTARY DATA

Supplementary Figure S1. Sequence conservation between GAAPs and BsYetJ.

Supplementary Figure S2. Structural models of AtGAAP1 define a putative ion channel pore.

Supplementary Figure S3. Structural models of AtGAAP2 define a putative ion channel pore.

Supplementary Figure S4. Structural models of AtGAAP4 define a putative ion channel pore.

Supplementary Figure S5. Structural models of AtGAAP5 define a putative ion channel pore.

Supplementary Figure S6. Structural models of AtGAAPs define a putative ion channel pore.

Supplementary Figure S7. Fluorescently tagged AtGAAPs rescue yeast from Bax-mediated apoptosis.

Supplementary Figure S8. Histochemical analysis of GUS expression in *proAtGAAP1-5::uidA* transgenic plants.

Supplementary Figure S9. *AtGAAP* gene expression analysis.

Supplementary Figure S10. RT-PCR analysis of *AtGAAP1-5* gene expression in the wild-type and *atgaap* mutant plants.

Supplementary Figure S11. Phenotypes of *atgaap* single, double and triple mutants.

Supplementary Figure S12. Subcellular localisation of AtGAAP1-YFP fusion protein in transgenic Arabidopsis lines.

Supplementary Table S1. Designation of Arabidopsis GAAPs.

Supplementary Table S2. Comparison of deduced amino acid sequences of human GAAP, viral GAAP and Arabidopsis GAAPs.

## ACKNOWLEDGEMENTS

Tuomas Puukko and Anna Huusari are acknowledged for excellent technical assistance. We thank Mikael Brosché for comments on the manuscript. This work was supported by the Biotechnology and Biological Sciences Research Council (BBSRC) (M.S., D.L.P, C.W.T and B.F.), the Wellcome Trust (C.W.T. and 090315 to G.L.S.), the Academy of Finland (275632, 283139, and 312498 to M.W.), the United Kingdom Medical Research Council (MRC) (G0900224 to G.L.S.), the University of Helsinki (M.W.), a Meres research associateship from St. John’s College, Cambridge (D.L.P.), the Isaac Newton Trust (G.C.), the Ella and Georg Ehrnrooth Foundation (M.S.), the Finnish Cultural Foundation (M.S.) and the Portuguese Foundation for Science and Technology - FCT (UID/DTP/04567/2016 and SFRH/BD/40714/2007 to N.S.). G.L.S. is a Wellcome Trust Principal Research Fellow. M.S., A.V. and M.W. are members of the Centre of Excellence in Molecular Biology of Primary Producers (2014-2019) funded by the Academy of Finland.

